# Range and elevation predict responses to climate change in frogs and lizards in the Western Ghats biodiversity hotspot of peninsular India

**DOI:** 10.1101/2025.04.10.648234

**Authors:** Aniruddha Marathe, S.P. Vijayakumar, Varun Torsekar, K.P. Dinesh, Saunak Pal, Achyuthan Srikanthan, Kartik Shanker

## Abstract

Anthropogenic climate change is altering the environment at unprecedented rates with severe consequences for most living organisms. As a result, species may extend, truncate, or shift their ranges in order to adapt to changing conditions. Species Distribution Models (SDMs) provide a data driven approach to predict future distributions under climate change and prioritize areas for conservation. Here, using a 10-year dataset with 4049 occurrence records of frogs from eight families and 27 genera as well as 733 occurrences of lizards from two families and 11 genera across the Western Ghats, we built SDMs to assess the changes in species distributions due to climate change. As expected, the temperature gradient across elevation and seasonality gradient across latitude contribute most to the climatic limits of species distributions. Latitudinal extents of most species were narrower in future predictions compared to the present, but there was little shift in latitudinal positions. On the other hand, most species shifted their distributions towards higher elevations, but the elevational range sizes remained the same. A total of 75 species of frogs (55%) and 15 species of lizards (45%) lose more than half of the suitable area, with few exceptions in both taxa that show an increase. In cases where a shift or increase in distribution was observed, the ability of the species to access and survive in these areas remains uncertain due to discontinuous topography and the presence of sister species. Overall, the frog and lizard fauna of the Western Ghats will be severely affected by climate change in the future due to a loss in climatic suitability.

## Introduction

Anthropogenic climate change is altering the environment at unprecedented rates with severe consequences for most living organisms (Walther *et al*. 2002, Malcolm *et al*. 2006, Parmesan 2006). Changes in the environment can potentially cause mismatches between species adaptations and environmental conditions. The response of species to novel environmental conditions can change species distributions towards new potentially suitable areas. As a result, species may extend, truncate, or shift their ranges (Shoo *et al*. 2006, Colwell *et al*. 2008, Colwell & Rangel 2010, Warren & Chick 2013). Additionally, natural and anthropogenic barriers to dispersal can reduce opportunities for species to track environmental changes, ultimately causing extinctions (Boulangeat *et al*. 2012, Schloss *et al*. 2012). Therefore, understanding the impacts of climate change on species distributions has become a critical conservation challenge.

As global temperatures rise, species ranges are expected to shift along latitudes and elevation in order to track the suitable climates (Chen, Hill, Ohlemüller, *et al*. 2011, Jacobsen 2020). Overall, the outcome of changes in species distributions due to climate change will be determined by the ability of a species to survive the changing environment and disperse towards suitable areas (Lenoir & Svenning 2015). Observations of latitudinal shifts due to climate change are largely from temperate rather than tropical latitudes (Sheldon 2019), including latitudinal shifts in the distributions of birds (Zuckerberg *et al*. 2009, Brommer *et al*. 2012) and butterflies (Parmesan *et al*. 1999) among other taxa (Hickling *et al*. 2006). On the other hand, tropical species are expected to respond by adjusting elevational distributions. This is because the latitudinal temperature gradient in the tropics is comparatively flatter (Sheldon 2019) and much of the change in climate takes place along elevational gradients (Colwell *et al*. 2008). For instance, frogs in the North Americas are expected to shift poleward, whereas those in the tropical South Americas are expected to shift to higher elevations (Lawler *et al*. 2009, Blaustein *et al*. 2010). Comparison of current and past distributions of trees (Feeley *et al*. 2011) and birds (Dejean *et al*. 2011, Forero-Medina, Terborgh, *et al*. 2011) on the Andean Mountains show evidence of upward shifts in species distribution. Model predictions from other regions for diverse taxa, including invertebrates (Chen, Hill, Shiu, *et al*. 2011), frogs (Forero-Medina, Joppa, *et al*. 2011), lizards (Jiang *et al*. 2023), birds (Sekercioglu *et al*. 2008) and mammals (Moritz *et al*. 2008, Rowe & Terry 2014), also support shifts in distributions towards higher elevations. Further, on mountain regions that span large latitudinal extents, such as the Andean Mountain range in South America, and the Western Ghats escarpment in India, latitudinal extents of species can reduce due to extinctions from local elevational gradients.

This is particularly important for the species in the Western Ghats (WG) – a region of high endemism and of global importance for conservation – due to the combined effects of seasonality gradients, and discontinuous topography of the mountain range. The Western Ghats (WG) harbors rich biodiversity, including numerous endemic species, and provide valuable ecosystem services (Das *et al*. 2006, Gunawardene *et al*. 2007). Climatic gradients across the latitudinal and elevational extents of the mountain range are important determinants of species distribution and community composition (Page & Shanker 2018, Bose *et al*. 2019, Jins *et al*. 2021). These gradients are likely to be altered in the near future due to climate change (Kumar *et al*. 2006). In addition, natural barriers in the mountain range and habitat loss in recent decades is likely to prevent species from colonizing suitable habitats, thus amplifying the impacts of climate change (Subba *et al*. 2018). Therefore, understanding potential species distributions and resulting community composition under climate change becomes crucial for biodiversity conservation in the Western Ghats.

The Western Ghats escarpment harbours a large number of endemic radiations of amphibians and reptiles due to a combination of isolation during the geological past, historical climatic fluctuations, and natural breaks in the mountain range (Vijayakumar *et al*. 2016, Mallik *et al*. 2020). As a result, there are numerous species with narrow geographic extents and environmental tolerances. Such species are particularly vulnerable to climate change as they are more likely to experience environmental mismatches in the future. Further, geographic barriers can prevent them from tracking suitable landscapes in the future, ultimately accelerating extinctions. Studies on climate change impacts in the Western Ghats on species of conservation focus (Priti *et al*. 2016, Sony *et al*. 2018), agriculturally important organisms (Sen, Gode, *et al*. 2016, Pramanik *et al*. 2021), and invertebrates (Sen, Shivaprakash, *et al*. 2016) suggest that the species distributions will be considerably altered. Still, what geographic areas, or which range attributes are vulnerable remains unknown as assessments across taxa have not been carried out.

We used a framework which uses geometric constraints in both latitudinal and elevational domains to make predictions regarding changes in range size and position (Colwell et al 2008). According to this framework, range sizes and range locations (midpoints) are related through geometric constraints, and reduction in suitable area due to elevational range shifts should be dependent on the range location. Therefore, there should be an interaction between current elevational extents and elevational midpoints. The total land area decreases towards higher elevations globally, however at landscape scales the relation between area and elevation can be variable (Elsen & Tingley 2015). Within the Western Ghats, land area decreases with elevation above 500m which is 80% of the total area of the escarpment. Therefore, we expect that the species shifting towards higher elevations should show decrease in suitable area.

Here, we use predictions from Species Distribution models (SDM) with climate in the present and future as a proxy for range extents of species to provide an assessment of the vulnerability of frog and lizard species to climate change in the Western Ghats. Our primary questions include: (1) What is the extent of change in climatic suitability for frogs and lizards in the Western Ghats? (2) How do attributes of species range such as latitudinal as well as elevational extents and location (midpoints) change under future conditions? (3) What range attributes explain the loss or gain in climatic suitability? (4) What consequences does this have for the frogs and lizards of the Western Ghats?

## Methods

### Occurrence data

We used occurrence records from extensive surveys across the Western Ghats from 2009 to 2016 as presence locations for the distribution models. The surveys covered topographic, elevational, latitudinal and rainfall gradients and were designed to capture latitudinal as well as elevational extents of frog and lizard species within the Western Ghats. The final dataset contains 4049 occurrence records across 203 species of frogs, and 733 occurrences across 59 species of lizards. Among these, we first removed any occurrences for the same species that were present on the same raster cell of the predictor variables, and further only considered species with five or more occurrence records. These criteria resulted in a set of 133 species with 3226 total occurrence records of frogs, and 32 species with 558 total occurrence records of lizards. All known families of frogs except Nasikabatrachidae which is represented by a single genus *Nasikabatrachus*, are present in the data, while lizard data comprised species in Agamidae and Gekkonidae only (and do not include Varanidae, Scincidae and Lacertidae).

### Environmental data

We initially considered 19 ‘bioclim’ variables available from Worldclim (Hijmans et al. 2005), and aridity index (Trabucco & Zomer n.d.) as climatic predictors, and finally used a subset of the layers after accounting for collinearity. The resolution of climatic variables is 30 arc sec (approximately 1 km^2^). Topographic information was included through elevation, slope, and aspect derived from the ASTER DEM at 30m resolution (Abrams et al. 2020). Topographic variables were re-sampled to match the coarser resolution of climate data. All the raster layers were clipped to the geographic extent (binding box) of the Western Ghats. We checked for cross correlations among all possible pairs of predictor variables using Pearson’s correlation coefficient and removed one predictor from the pairs that had correlation coefficient equal to or more extreme than ±0.7. The final subset represents variation in annual mean values as well as seasonality of temperature and precipitation (Table 1). We used these variables as predictors for all the species as a reduced subset of environmental variables for mean and annual variability in rainfall and temperature is recommended for modeling distributions of large numbers of species with limited ecological information (Low et. al. 2021).

**Table 1.** predictor variable used in distribution models.

| Predictor | Description |
| --- | --- |
| Bioclimatic |  |
| Bio 1 | Annual Mean Temperature |
| Bio 4 | Standard deviation of temperature x 100 |
| Bio 12 | Annual Precipitation |
| Bio 14 | Precipitation of Driest Month |
| Bio 15 | Precipitation Seasonality (Coefficient of Variation) |
| Bio 18 | Precipitation of Warmest Quarter |
| Bio 19 | Precipitation of Coldest Quarter |

### Distribution modeling

We predicted geographic distributions of species based on climatic suitability using the MAXENT algorithm (Philips *et. al.* 2006). As it is nearly impossible to infer true absences from survey records alone, we used the species occurrence records as presence only data. The MAXENT algorithm contrasts distribution of environmental variables at occurrence locations against environmental conditions in the entire region to generate predictions of suitability. The observed distributions can be potentially biased due to unequal or incomplete sampling. While the sampling effort was kept as comparable as possible across the latitudinal extents, inherent differences in habitat availability and geography of the Western Ghats result in greater number of occurrence records towards lower latitudes. To account for this difference, we generated a bias layer which is an inverse distance weighted density kernel, based on pairwise distances between occurrence points, using the command ‘kde2d’ in the package ‘MASS v7.3’. The resolution and extent of the bias file was set to match the predictor variables. From the bias raster, 10,000 cell centroids were randomly sampled weighted by the value for each cell. Environment represented by these points represents any geographic bias that may be present in the samples. These points were used as biased background locations in MAXENT. We constructed separate bias layers for the different families of frogs and lizards.

We compare several maxent models with different combinations of feature classes, and regularization parameters to select an optimum model by running the maxent algorithm through the R package ENMEval v2.0 (). The package compares models with different combinations of feature classes, and regularization parameters, and estimates model evaluation statistics. We used linear, quadratic, and hinge feature classes, and their combinations, as linear+quadratic, linear+hinge,quadratic+hinge, and linear+quadratic+hinge, with a range of regularization parameters from 0.5 to 5 at an interval of 0.5. Model cross validation was done using the Spatial block method with default parameters for species with more than 10 occurrences and with Jackknife method for species with 10 or fewer occurrences. Final model among the candidate models was selected based on sequential criteria of (1) Training Area Under Curve >0.6 (2) minimum difference between training and testing AUC (3) minimum number of coefficients (4) maximum value for regularization parameter (5) minimum number of feature classes. The AUC (Area under receiver operator Curve) was used to evaluate model performance. AUC is analogous to ranked correlation of predicted probability between presence and absence points. High correlation suggests the model can effectively distinguish between potentially suitable areas from the background. All continuous predictions were converted to binary (suitable/unsuitable) using the ‘Maximum sum of sensitivity and specificity’ threshold criterion (Liu *et al*. 2005)

In addition, we also built models with an unbiased background using the same predictors and model parameters. We visually examined the model predictions with and without bias. Accounting for bias appeared to have minimal apparent effect on model predictions. However, for a small number of species, typically when the occurrence points did not coincide with occurrence density of the family, predictions with a biased background extended far beyond the latitudinal extents of occurrence records. Potential overpredictions were confirmed by consulting taxon experts who had participated in the survey effort to generate the occurrence data and have extensive taxonomic and ecological knowledge of these groups. Following this consultation, predictions with unbiased background points were used for some species (Supplementary data). In addition, we modeled distributions using Ensemble models implemented in using R package biomod2 v3.4 visual inspections by experts suggested that these models performed at best as well as MAXENT. Hence, we used only MAXENT for all predictions as it uses a single algorithm and is easier to interpret.

The distribution models were projected to future climatic scenarios of SSP1-2.6, and SSP5-8.5 for the period 2061-2080. For selecting the climate models that will be used for future projection, we identified the most pessimistic and optimistic conditions for the Western Ghats, by comparing trends in the projected temperature values across the Western Ghats for all models available in the ‘Worldclim’ repository and ranked the models based on predicted increase in temperature.We used three models with least projected increase in temperature for representing the optimistic SSP1-2.6 scenario and three models with the most increase in temperature for representing the pessimistic SSP5-8.5 scenario (See supplementary-1 for details). We predicted the species distributions using the three selected models in each scenario and used the mean of the predictions as the predicted distribution of the species under each scenario.

### Identifying predictions beyond biogeographic limits

The Western Ghats show marked climatic changes which generate a gradient of increasing seasonality towards higher latitudes. These changes very likely limit species distributions in the region and in turn contribute to the model predictions. In addition, the species distributions are limited by the discontinuity in the mountain range, such that species are unable to access suitable climates separated across valleys. Due to evolutionary processes, phylogenetically and ecologically related species may occupy similar environmental conditions on either side of such barriers. For a larger proportion of frogs and lizards in the WG, sister species are distributed across topographic barriers, likely a consequence of allopatric speciation events; these sister species are often morphologically similar and occupy similar habitats (Vijayakumar *et al*. 2016, Robin *et al*. 2017, Mallik *et al*. 2020)

However, the distribution models that are predicting climatic suitability do not directly account for these processes and should predict suitability in areas that are either inaccessible to the species or areas that overlap the distributions of sister species. This can result in overprediction of geographic distribution. Thus, to reduce overpredictions in areas that are disconnected due to geographic features in the mountain range, we identified the subset of the predicted distribution for every species that is most likely accessible to the species, using a combination of criteria based on the known distribution, elevational extents, and elevation of the discontinuous contour.

We used information on the known distributions of phylogenetically related species of bush frogs (Rhacophoridae), and torrent frogs (Nyctibatrachidae) to identify 17 massifs that are most likely to be dispersal barriers for frogs in the Western Ghats. We then mapped the lowest contours that form discontinuous elevations between these massifs. We considered all massifs within latitudinal limits of the observed distribution as potentially accessible areas. Then, we identified suitable areas predicted in MAXENT that were present within these limits. Further, when sufficient phylogenetic data was available, we confirmed the overpredictions using occurrences of phylogenetically related species. The resulting predictions combine the effects of climatic constraints as well as biogeographic processes in the Western Ghats and provide a more nuanced approach towards predicting habitat/climatic suitability in the present or future. We report the suitable areas both within and across barriers for each species.

### Identification of high diversity areas

Following Srinivasulu et. al. (2021), we mapped areas containing high diversity within the Western Ghats by calculating the mean of the binary prediction for all species and rescaling the resulting raster from 0 to 1. Values of 0.5 or higher represent areas that have at least half the species compared to the maximum species richness.

### Latitudinal and elevational extents of species

The latitudinal extent of the entire predicted distribution is an overestimation of the species actual latitudinal extents in most cases. This is because small numbers of disconnected pixels may be predicted much beyond the extents where a majority of the suitability is predicted. To reduce potential overpredictions, we excluded 5% of the area towards the northernmost and southernmost extents and the remaining 90% of the prediction was considered for estimating latitudinal extents of each species. To obtain these estimates, we first divided the WG into 0.8^0^ latitudinal zones, and calculated the fraction of total predicted distribution within each latitudinal zone. The northern and southern extreme latitudes that contained 90% of the predicted area were used as latitudinal extent of the species.

We estimated the elevational extents of species within each 0.8^0^ degree latitudinal zone separately in the same manner by using 200m elevation zones, such that each species had separate elevational extents in each latitudinal zone. Further, we estimated elevational midpoints for each species as the mean of midpoints on all massifs weighted by the suitable area for the species in the massif. We compared the latitudinal and elevational extents of species in present and future predictions and confirmed the patterns using linear models.

### Predictors of changes in suitable area under climate change

Species that occur across small geographic extents or are not adapted to withstand environmental variability are more likely to go extinct due to climate change. We tested this using the latitudinal and elevational extents and midpoints under current conditions and the decrease in suitability in future predictions. We tested if there is any difference between the range attributes i.e midpoints and extents of species that show near complete loss of suitable area in the future predictions and the entire species pool. For this, we considered species for which predicted area in future is <1% of the current prediction. We obtained the distributions of current latitudinal/elevational extents and midpoints for these species and compared these with the regional species pool. Standardized Effect Sizes (SES) were calculated with 1000 bootstrap randomizations.

We also used linear regression models to test the relationship between change in area and attributes of current as well as future predicted range extents; we used current latitudinal and elevational extents and midpoints along with difference in the midpoints between present and future as predictors for the regression models. To test these predictions, we used a linear regression model with all the possible predictors and simplified it based on AICc. Further, for the frog data, we demonstrate the effects of difference in elevational midpoints and the interaction term by reporting change in AICc and R^2^. In the case of lizards, as there are only 32 species in the data, we only used predictors relating to elevational ranges.

## Results

Based on the data availability criteria, 133 species of frogs that had at least five occurrence records each located on separate raster cells were selected for distribution modeling. The AUC values of distribution models ranged between 0.6 and 0.99. However, 80% of the species had AUC values greater than 0.8. For lizards, 32 species were used for distribution modeling. The AUC values ranged 0.62 and 0.99. Mean annual temperature and dry period precipitation had the highest percentage contribution for most species (38 species of frogs, 10 species of lizards) followed by precipitation seasonality (24 species of frogs, and 6 species of lizards).

A large proportion of species (92 out of 133 species) were recorded from up to four massifs out of the 16. Only two species - *Duttaphrynus melanostictus* and *Pedostibes tuberculosus* were distributed across most of the massifs (14). There are 33 species, mostly from the families Rhacophoridae and Nyctibatrachidae, that have occurrences on a single massif. All species of lizards except for *Agasthyagama bedommei* occurred on more than one massif and for 7 species, 70% of the predicted area was on disconnected massifs.

For several species the suitable area identified as beyond elevational barriers, was across the Palaghat or Shencottah gap, which are elevational barriers in the escarpment that separate allopatric species of several taxa. In other cases a large proportion of area of the predicted distribution overlapped with a phylogenetically related sister species (eg *Nyctibatrachus sanctipalustris and Nyctibatrachus karnatakensis; Indosylvirana intermediatus and Indosylvirana caesari*).

### Predictions for future climates

Suitable area predicted by distribution models decreased for most species (Table 2; Figure 1). At least 19 species of frogs may potentially go extinct as suitable areas in future are less than 1% of the current predictions. Most of these species belong to the endemic genera *Nyctibatrachus* and *Raorchestes*. Among other species, the median of the suitable area predicted was 50% of the current distribution. In the case of lizards, only two species, namely *Cnemaspis indica* and *Agastyagama beddomei*, show nearly complete loss of suitable area in the future, while *Cnemaspis belarion* (80% - 99% decrease) and *Cnemaspis galaxia* (15% - 98% decrease) lose a majority of their suitable area.

**Figure 1.**
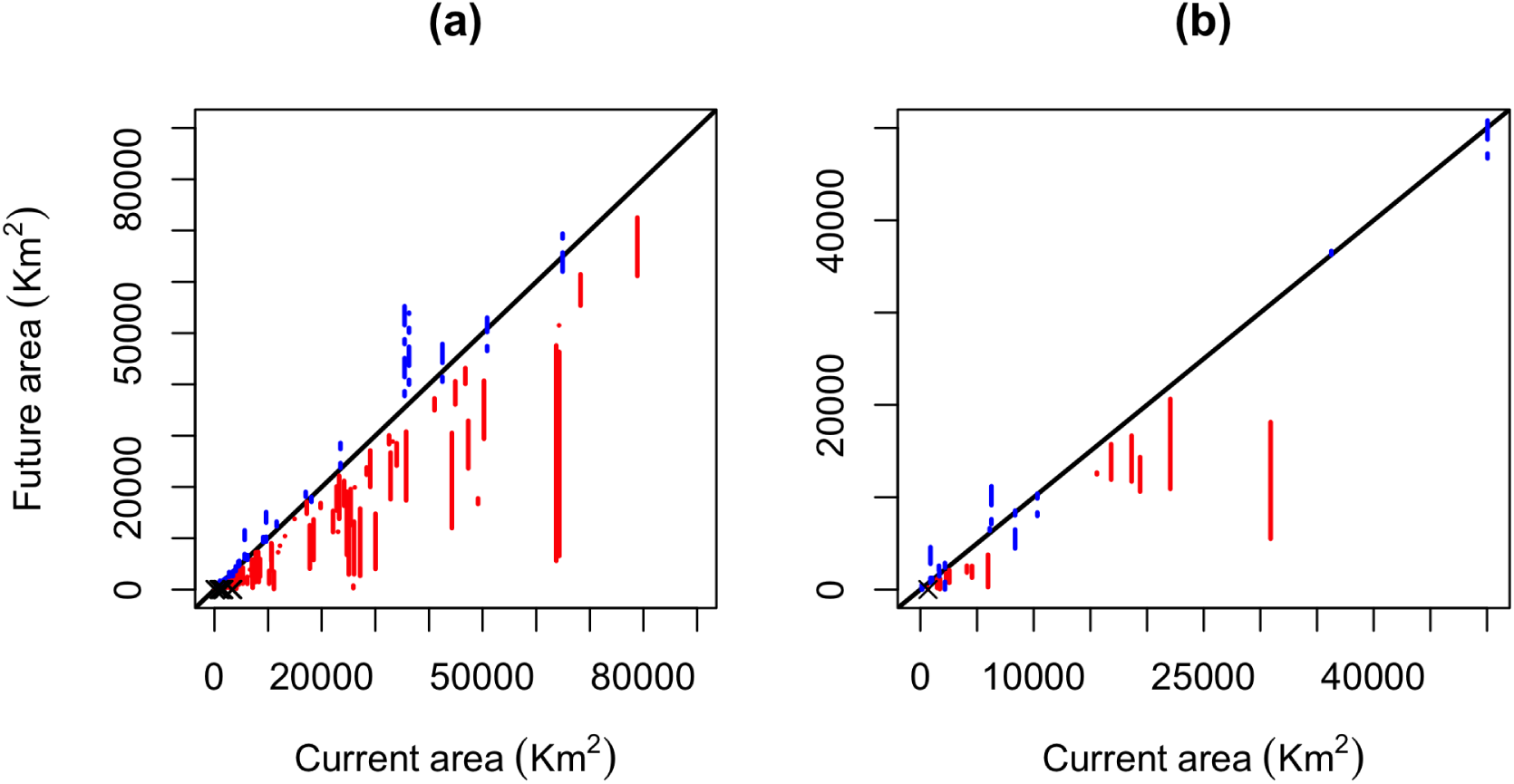
Suitable areas predicted under climate change compared to currently suitable area for frogs (a) and for lizards (b). Variation expected due to differences among scenarios is represented as vertical segments. Solid segments show reduction in area while dotted segments show increase in suitable area in the future.

**Table 2.** Percentage of species within each family for which suitable area decreased or increased in the future climate scenarios.

|  |  | SSP1-2.6 |  | SSP5-8.5 |  |
| --- | --- | --- | --- | --- | --- |
|  |  | decrease | increase | decrease | increase |
| Frogs | Bufonidae | 83.3 | 16.7 | 66.6 | 33.4 |
|  | Dicroglossidae | 100 |  | 100 |  |
|  | Micrixalidae | 81.8 | 18.2 | 100 |  |
|  | Microhylidae | 60 | 40 | 70 | 30 |
|  | Nyctibatrachidae | 78.5 | 21.5 | 85.7 | 14.3 |
|  | Ranidae | 84.6 | 15.4 | 84.6 | 15.4 |
|  | Ranixalidae | 66.6 | 33.4 | 66.6 | 33.4 |
|  | Rhacophoridae | 80.3 | 19.7 | 96 | 4 |
| Lizards | Agamidae | 71.4 | 28.6 | 92.8 | 7.2 |
|  | Geckonidae | 72.2 | 27.8 | 61.1 | 38.9 |

As expected,the reduction in suitable area was less severe for the relatively optimistic conditions, but 10 species of frogs and three species of lizards showed greater suitable area under SSP5-8.5 compared to SSP1-2.6 contrary to the expectations. Forteen species of frogs show an increase in suitable area even under the most pessimistic projections. In most areas of increase are spatially disconnected (eg. Nyctibatrachus urbis, and Nyctibatrachus shiradi) and present on a separate elevational contour.

### Latitudinal and elevational extents of species

Latitudinal extents were smaller in future compared to the present for frogs (w = 6023,p<0.05,Figure 2), but not for lizards (Figure 2,w =, p). Across all current latitudinal extents, the median of predicted extents in the future (SSP5-8.5) was consistently smaller (Figure 2). The patterns in latitudinal midpoints or range locations were less clear but statistically significant for frogs (Figure 2;w =, p) and not for lizards (Figure 2; wil).

**Figure 2.**
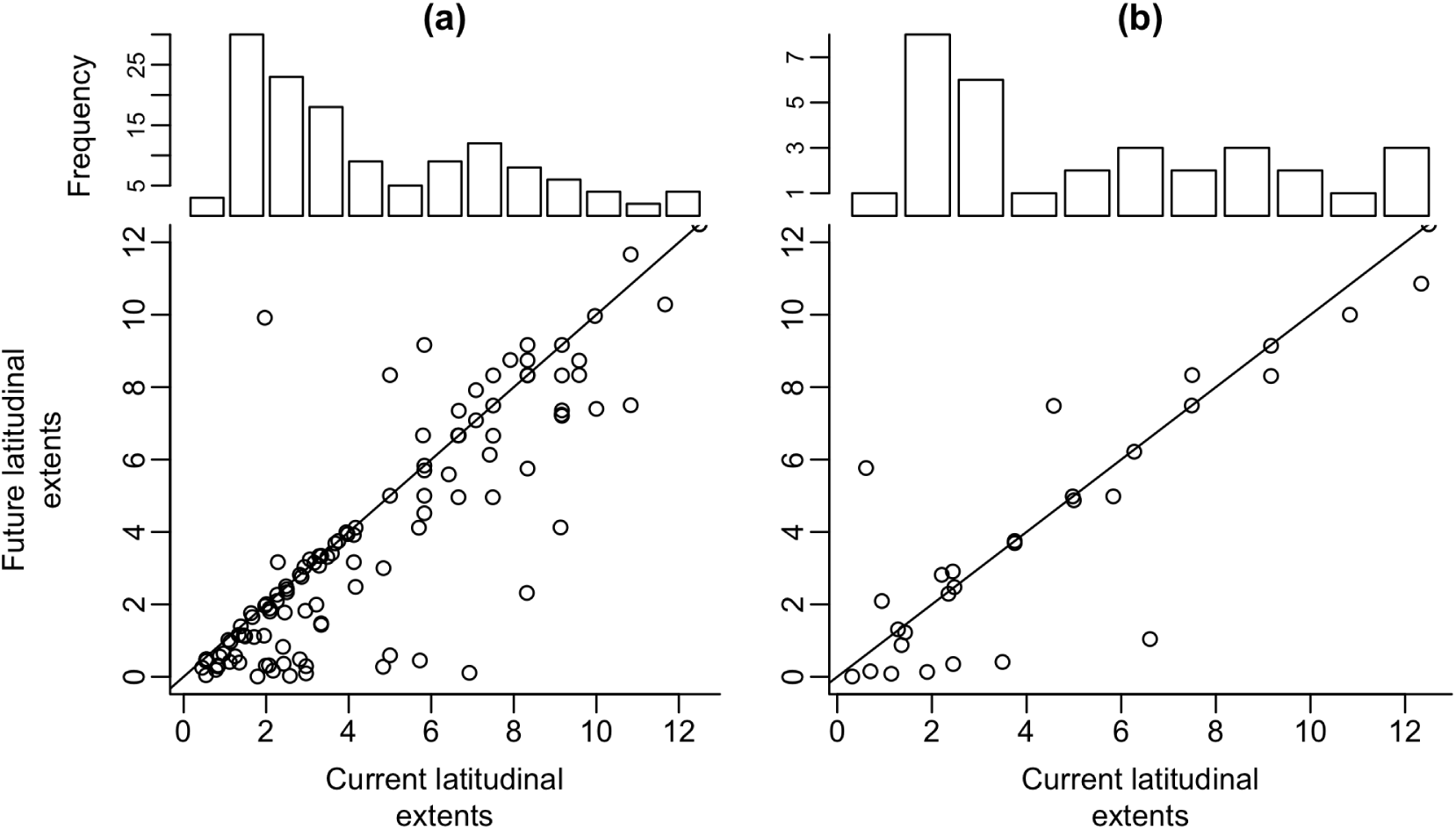
Latitudinal range extents of frogs (a) and lizards(b) in future (SSP5-8.5) compared to the present. The Histograms in top panels show the number of species that have the corresponding range extents.

Elevational extents (Figure 3) of frogs were smaller in future compared to the present (w = 9488, p<0.05) but not for lizards (w = p = 0.3). On the other hand, Elevational midpoints were higher for both frogs and lizards (Figure 4; Frogs: w = 6355, p<0.05; Lizards: w =, p<0.05).

**Figure 3.**
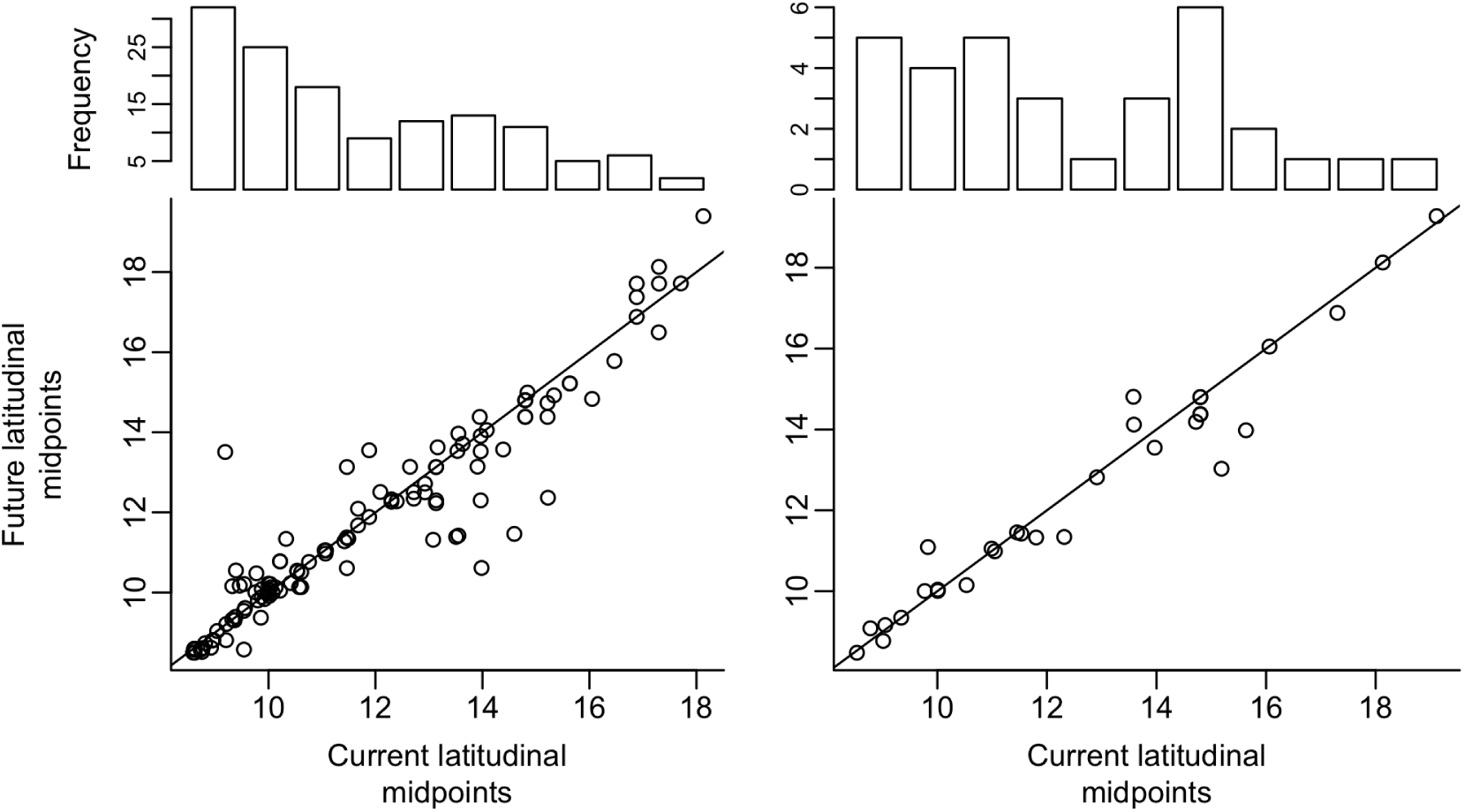
Latitudinal midpoints of frogs (a) and lizards(b) in future (SSP5-8.5) compared to the present. The Histograms in top panels show the number of species that have the corresponding range extents.

**Figure 4.**
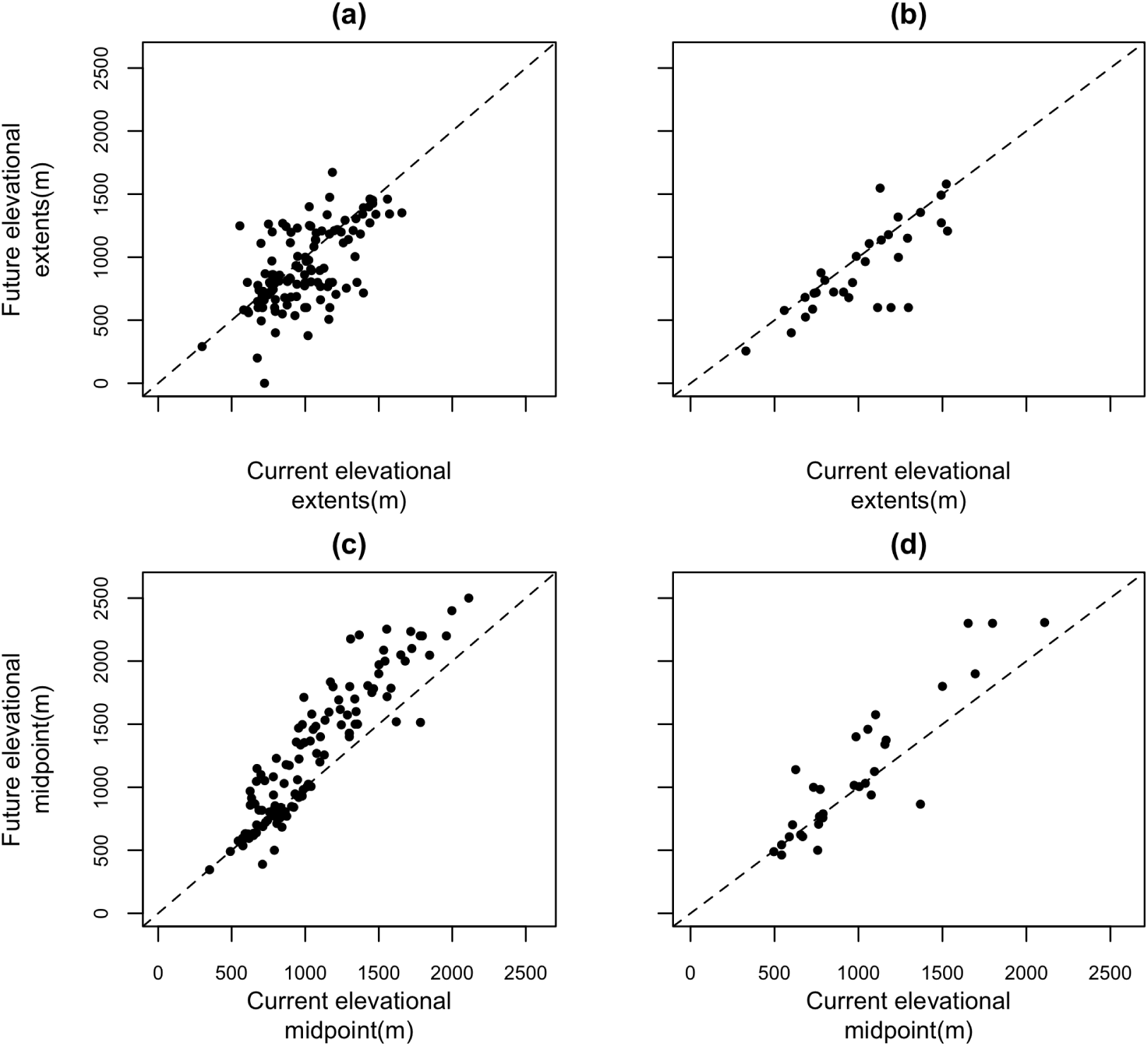
Elevational extents and midpoints of frogs (a, c) and of lizards (b, d) in future (SSP5-8.5) compared to the present. Dashed lines are 1:1 lines indicating no change.

### Areas of high diversity under present and future climates

Based on the stacked binary distribution models, approximately 5% of the total area of the WG for frogs, and 8% for lizards, was found to be suitable for at least half the species compared to the maximum reported species richness (Figure 5). Most of these areas were in the Southern Western Ghats, particularly the Agasthyamalai mountain (8^0^ N to 8.9^0^N, and the peaks of the Anamalai and Meghamalai mountains above 1000m elevation (8.9^0^ to 10.3^0^N). North of the Palghat gap, the high diversity areas mark the 600m contour with the exception of the Nilgiri plateau (11.0^0^N) at more than 1000m elevation, and the Castle rock region (14.5^0^N – 15.7^0^N) forming a low elevation plateau north of the Agumbe range (14.0^0^N)

**Figure 5.**
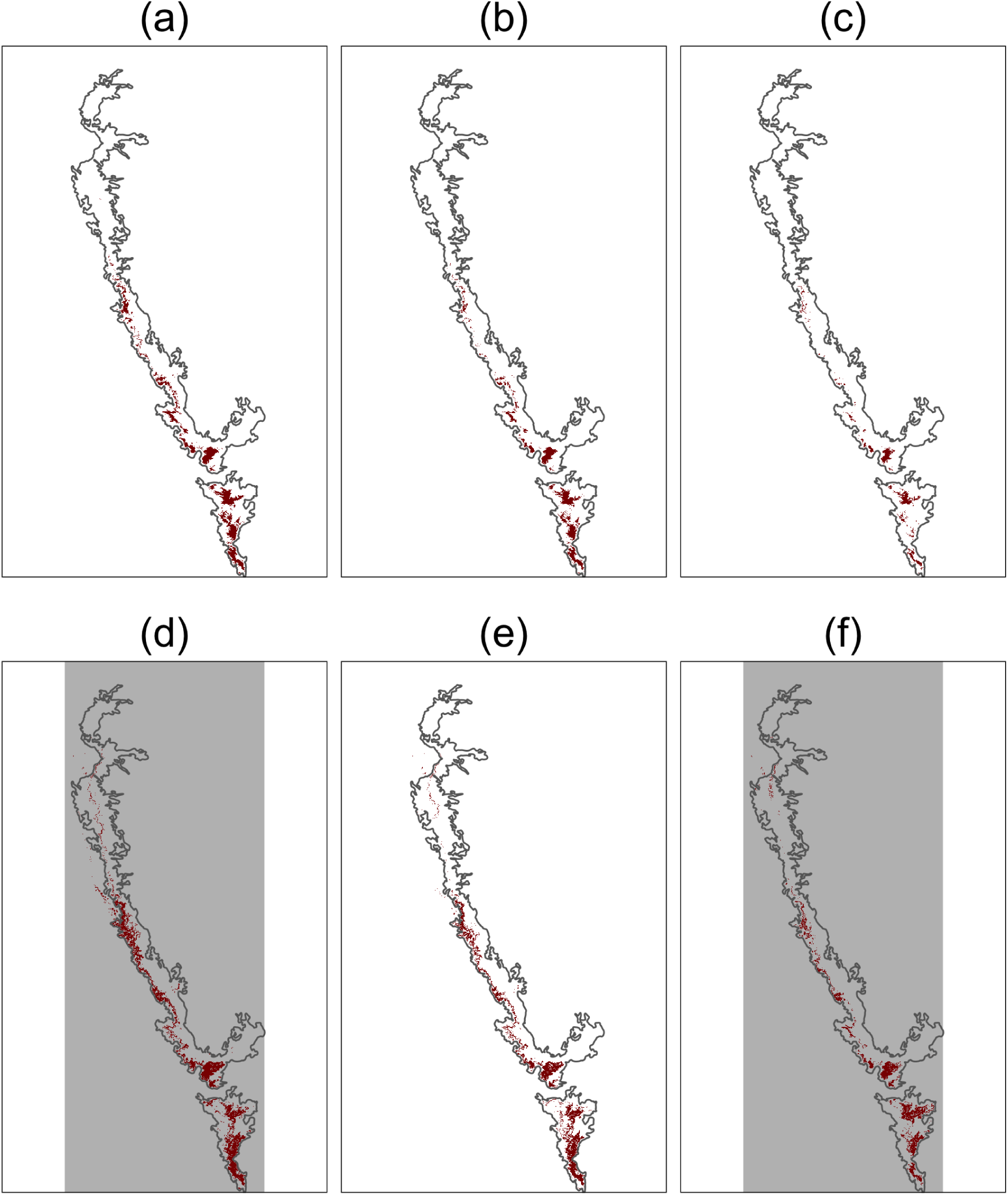
Areas with High diversity of frogs (a,b,c) and for lizards (d,e,f) within the Western Ghats under current (a,d), SSP1-2.6 (b,e) and SSP5-8.5 (c,f) scenarios and for lizards

Under climate change, the high diversity areas reduced to less than 2% for SSP1-2.6, and 0.4% for SSP5-8.5 scenarios. The areas predicted with future scenarios were subsets of the area predicted under the current conditions and major shifts towards higher or lower latitudes were not observed. The predicted high diversity areas were typically restricted within higher elevation contours with loss of suitability in lower elevations compared to the present.

### Predictors of change in area

Bootstrap analysis showed that species that are most likely to go extinct in the future have narrower latitudinal (SES = -3.6, p<0.01) as well as elevational extents (SES = -1.7, p<0.05) and lower latitudinal midpoints (SES = -2.2,p<0.05) and higher elevational midpoints (SES = 2.81) (Figure 6). Latitudinal extents and midpoints were highly cross correlated (pearson’s correlation = 0.8). Therefore, we only included latitudinal extents in the model and did not include the interaction between the two predictors in the full model. The Regression model for percentage of suitable area in future for frogs with all predictors explained 42% of variance. Dropping latitudinal extent from the model did reduce the AICc. The final model contains negative effects of elevational location, as well as upward shift in elevation. The interaction effect between elevational range and location is weak and it indicates that the effect of elevational location is marginally smaller at larger elevational extents. On the other hand, exploratory analysis of interaction between latitudinal extents and midpoints, showed a positive effect of latitudinal extents, at lower and intermediate latitudes but not at higher latitude.

**Figure 6.**
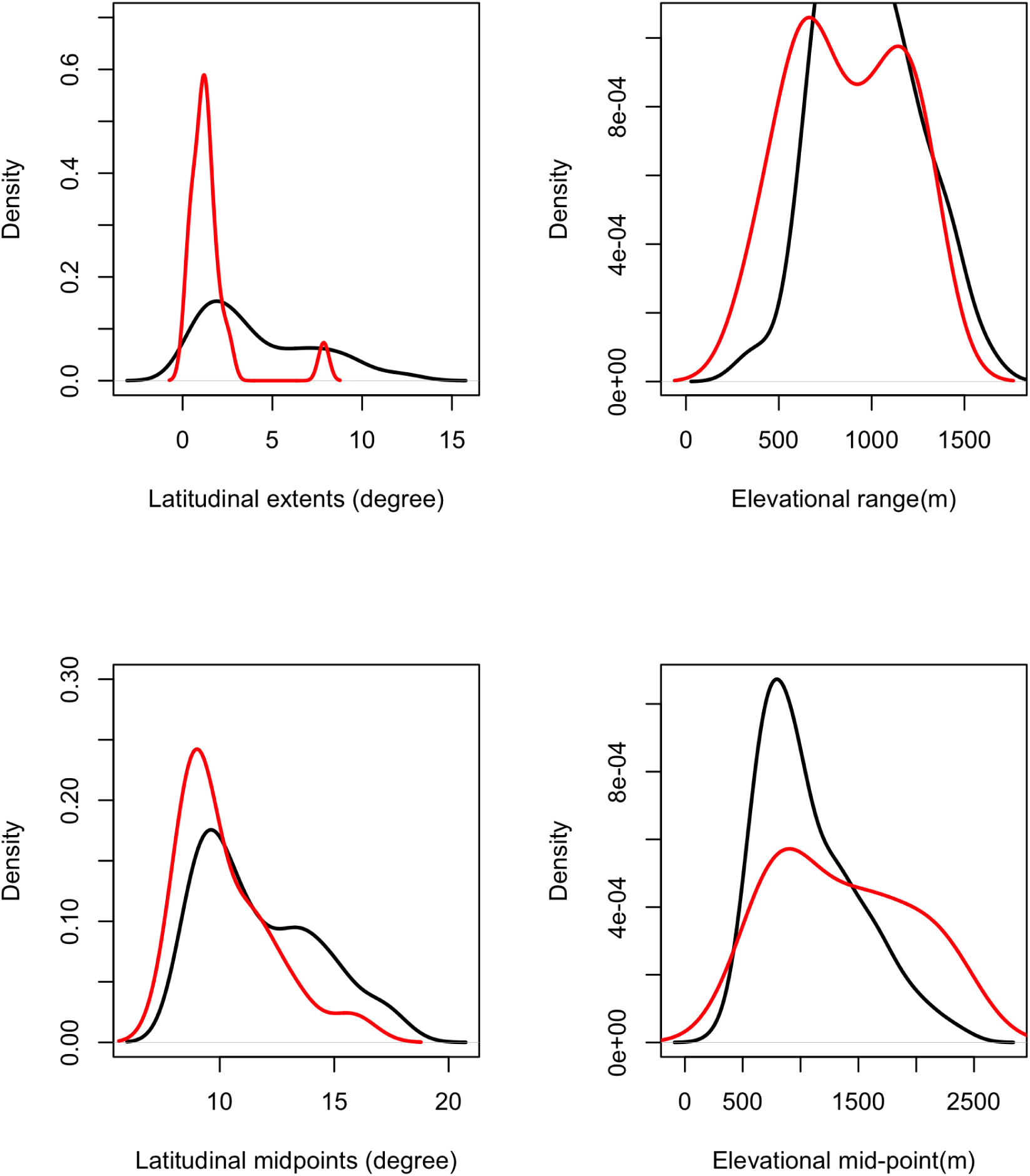
Density kernel for elevational midpoints (a) latitudinal extents (b) elevational extents, (c) latitudinal midpoints, and (d) elevational midpoints of entire species pool (black lines) and the species have suitable area in SSP5-8.5 which less than 1% of current suitability (red lines).

In the case for lizards,the model contains positive effect of elevational range and negative effect of elevational position and elevational shift, along with interaction between elevational extents and midpoints. The proportion of area in the future projection, increases with elevational extent at low elevations but the relation becomes weaker towards higher elevation.

## Discussion

The Western Ghats Escarpment is a region of global importance for the conservation of biodiversity due to a large number of endemic species. Therefore, estimating the effects of anthropogenic climate change on the biodiversity of the Western Ghats is of critical importance for conservation. Our results show that a combination of shifts in climatic suitability and geographic limits to species distribution will have adverse effects on the frog and reptile fauna of the mountain range. While studies do confirm the negative impact of anthropogenic climate change on plants (Sen, Gode, *et al*. 2016), invertebrates (Sen, Shivaprakash, *et al*. 2016) and mammals (Sony *et al*. 2018), assessments for species groups representing a large assemblage of species are still rare, with the notable recent exception of reptiles (Srinivasulu *et al*. 2021). Here, we present an assessment of the effects of climate change on the geographic distributions of frogs (133 species) and lizards (32 species) in the Western Ghats using the most comprehensive distribution data assembled to date for these taxa in the region.

Predicted suitable areas for most species decreased under climate change as shown in earlier studies (Sen, Gode, *et al*. 2016, Sony *et al*. 2018, Pramanik *et al*. 2021, Srinivasulu *et al*. 2021). As expected, most species displayed a large variation in predicted suitable area between the most optimistic and pessimistic scenarios. This confirms that the failure to arrest the rate of emissions and increase in temperature will have a severe negative impact on biodiversity. Of the 133 species, 35 species of frogs showed an increase in suitable area mostly on the periphery of current distribution, in the SSP1-2.6 scenario only. Further, for nine species, at least part of the increase in area is disconnected from the current distribution and so it is uncertain that these species can access these areas. This is particularly true for species with small latitudinal extents such as *Indosylvirana urbis, Nyctibatrachus deveni*, and *Nyctibatrachus karnatakaensis* but also the case for species with the largest increase in area - *Nyctibatrachus Shiradi*, and *Nyctibatrachus sylvaticus*. Though suitable areas might be present, it is unlikely that most of these species would be able to extend their ranges. On the other hand, some species with large latitudinal extents and large increase in suitability such as*, Euphlyctis karaavali*, and *Uperodon anamalaiensis* have disjunct distributions where only one or two occurrences are reported across a large gap in latitude. Future work may fill the gaps in distribution leading to different model predictions, or the species may be identified as cryptic species complexes, so that the suitability for different populations/species should be predicted separately. On the other hand, the increase in area for lizards takes place by extending the existing distribution around the existing suitable areas. For species that show a particularly large increase in area, for instance *Cnemaspis lithophilis* and *Sitana spinaecephalus*, the increase is not on spatially disconnected polygons where dispersal may be uncertain.

Within the Western Ghats, mean annual temperature shows less variation across latitude than across elevations (Page & Shanker 2020). Therefore, it is likely that this variable represents elevational limits of the species. On the other hand, seasonality in precipitation and temperature increases significantly towards higher latitudes (Page & Shanker 2020). Therefore, seasonality variables should contribute to the latitudinal limits of species distributions to a greater degree. As expected, mean annual temperature and precipitation seasonality were important variables for most of the species of frogs. On the other hand, temperature seasonality was found to be the most important variable, with minimal importance for precipitation seasonality, for lizards in this study and for squamates in general in other studies (Srinivasulu *et al*. 2021). Limits of water availability on amphibian life cycles very likely generate this difference.

High diversity areas show striking reduction with future climate projections. The changes reported here are drastic compared to similar results for reptiles. High diversity areas for reptiles covered a greater proportion of the Western Ghats, and remained largely contiguous even under the most pessimistic scenario in our study, as well as others (Srinivasulu *et al*. 2021). In comparison, the areas for amphibians are smaller and discontinuous even under current conditions. Further, under the effect of climate change, the high diversity areas shrink in size and remain restricted to higher elevations. This result confirms the prediction of ‘lowland biotic attrition’ (Colwell *et al*. 2008), i.e. low elevation assemblages lose species due to shifts in species distribution towards higher elevations. Elevational range shift is the most immediate response to climate change expected for tropical species (Colwell *et al*. 2008). Upward shifts in elevational distributions and extinction threat for high elevation montane species are also predicted for amphibian fauna of Madagascar (Raxworthy *et al*. 2008) and south-east Asia (Bickford *et al*. 2010). This is also supported in the present study as elevational distributions for frogs and lizards do shift towards higher elevation (Figure 4). Because mountains are naturally narrower near the summit, the amount of area available decreases towards higher elevations. Therefore, shifts in distributions towards higher elevations should decrease the total suitable area, other factors being constant. Shift in elevational midpoints is an important predictor of change in suitable areas for frogs as well as lizards, which supports the above prediction. As the space available for upward movement will reduce towards higher elevation, the effect of increase in elevational extents on future suitability should be greatest at low elevations and should reduce with increase in elevation. This should result in interaction between the effect of elevational extents and elevational midpoints. While the interaction is not a strong predictor of future suitability, exploratory analysis indicates that for frogs, the slopes of the relation are steeper at intermediate elevations and latitude.

The projected changes in suitability should be seen as conservative estimates of climate vulnerability, as biological mechanisms such as diseases, developmental errors due to accelerated metamorphosis, and decrease in dissolved oxygen in streams among others can accelerate the negative effects of climate change beyond changes in the climate envelope itself (Bickford *et al*. 2010, Blaustein *et al*. 2010, Li *et al*. 2013). The results presented here suggest that temperature gradients across elevation, and seasonality gradients across latitude, contribute most to the climatic limits of species distributions. Most species do not show latitudinal shifts in suitability as a response to climate change and in the few cases where such shifts are observed, the ability of the species to access and survive in these areas remains uncertain due to discontinuous topography and the presence of sister species. Overall, the results suggest that the herpetofauna of the Western Ghats will be severely affected by climate change in the near future, with greater impacts on frogs, in particular some endemic groups, than lizards.

**Table 3.** Linear regression models for change in area between current and SSP5-8.5 projections for frogs.

| Rank | Models | K | AICc | Delta AICc | AICc weight | R<sup>2</sup> |
| --- | --- | --- | --- | --- | --- | --- |
| 1 | elevational range + elevational mid-point + shift in elevational midpoint + shift in latitudinal midpoint | 6 | 1163.75 | 0 | 0.65 | 0.43 |
| 2 | elevational range + elevational mid-point + elevational range:elevational mid-point + shift in elevational midpoint + shift in latitudinal midpoint | 7 | 1165.5 | 1.81 | 0.26 | 0.43 |
| 3 | elevational range + elevational mid-point + elevational range:elevational mid-point + latitudinal range + shift in elevational midpoint + shift in latitudinal midpoint | 8 | 1167.64 | 3.88 | 0.09 | 0.43 |

**Table 4.** Coefficients of regression model for percentage of area in future predictions for frogs.

| Predictors | Estimate | Std. Error | p value |
| --- | --- | --- | --- |
| Intercept | 54.5 | 2.87 | <0.05 |
| Elevation extents | 5.76 | 3.04 | 0.06 |
| Elevation midpoints | -21.83 | 3.71 | <0.05 |
| Shift in elevation midpoints | -8.38 | 3.53 | 0.01 |
| shift in latitudinal midpoint | 10.13 | 3.12 | <0.05 |

**Table 5.** Coefficients of regression model for percentage of area in future predictions for lizards. Elevational and latitudinal predictors are part of two separate linear models. The response variable was log transformed.

| Predictors | Estimate | Std. Error | p value |
| --- | --- | --- | --- |
| Intercept | 3.67 | 0.22 | <0.05 |
| Elevation extents | 0.3 | 0.22 | 0.17 |
| Elevation midpoints | -0.2 | 0.3 | 0.5 |
| Shift in elevation midpoints | -0.63 | 0.22 | <0.01 |
| Elevation extents:Elevational midpoints | 0.65 | 0.28 | 0.05 |
| Latitudinal range | 0.46 | 0.24 | 0.06 |
| shift in latitudinal midpoint | 0.85 | 0.23 | 0.001 |

## Supporting information

Data for model parameters and range atributes of frogs

Data for model parameters and range atributes of lizards

Rscript file for producing figures and tables in the paper

Supplimentary file with descriptions of selecting climate models and exploratory analysis

## Acknowledgements

AM would like to thank the CSIR-Research Associate program for support during preparation of the manuscript and the state forest departments of Kerala, Karnataka, Tamil Nadu, Goa and Maharashtra for facilitating the fieldwork. VSP, VT, SP, and AS contributed the data for the analysis presented here. We acknowledge the support of the Director, ZSI, Kolkata and the Officer-in-Charge, ZSI, WRC, Pune.

AM would like to thank the CSIR-Research Associate program, and the IISC-DBT postdoctoral fellowship program for support during preparation of the manuscript and the state forest departments of Kerala, Karnataka, Tamil Nadu, Goa and Maharashtra for facilitating the fieldwork. VSP, VT, SP, and AS contributed the data for the analysis presented here. KPD acknowledges the support of the Director, ZSI, Kolkata and the Officer-in-Charge, ZSI, WRC, Pune for the support.

## Statements and Declarations

Funding : Critical Ecosystem Partnership Fund (CEPF) (), CSIR, DBT-IISc Partnership Programme (), Ministry of Environment, Forest and Climate Change (MOEFCC) (); Council of Scientific and Industrial Research (CSIR) – Research Associate fellowship ().

Permit: The following state forest departments provided permission for fieldwork and collection: Kerala, Karnataka, Tamil Nadu, Goa and Maharashtra.

Field team: Mrugank Prabhu, SR Chandramouli, Mayavan

### Author contributions

KS and AM conceptualized the study. AM performed the statistical analysis. VT, SPV and KPD collected the data, and provided key inputs for distribution models. AM wrote the first draft of the paper. KS edited the draft and provided inputs at all stages.

