## Supplementary material for "Range and elevation predict responses to climate change in frogs and lizards in the Western Ghats biodiversity hotspot of peninsular India": Supplimentary file with descriptions of selecting climate models and exploratory analysis

**Selecting climate models for future projections of SDMs**

The species distribution models can be projected using future climate scenarios to generate expected changes in species distributions under the impending climate change. The predictions for future climate conditions considering the optimistic and pessimistic scenarios of changes in carbon emission rates are made using several different models. Typically, changes in species distributions are reported for projections from multiple models to reduce the uncertainty in outputs. However, as the exact nature of the models and the differences among the models are not readily accessible to biologists, selecting the models using objective criteria becomes difficult. Here, we attempt to identify the most pessimistic and optimistic conditions for the western ghats, by comparing the projected temperature values across all models. We followed the following steps for selecting the climate models:

1. We considered the median values of mean annual temperature, within a 0.8 degree latitudinal zone of the western ghats predicted by all models available on worldclim.

2. We then identified the models that were predicting the lowest temperature for the optimistic scenario SSP1-2.6 and the highest temperature for the pessimistic scenario SSP5-8.5 at each latitude zone.

3. The models were then ranked based on the number of latitudinal zones in which the predicted temperature values are either highest or lowest. The first three models in both categories were selected. The selected models represent the extremes of the projected variation. Bioclim variables for the selected models were used to generate predictions of species distribution models under climate change.

Model selected for the SSP1-2.6 scenario are (1) "MIROC6", (2)"GFDL.ESM4", (3) "INM.CM5.0" and the three models for SSP5-8.5 are (1) "UKESM1.0.LL",(2) "ACCESS.CM2", (3) "HadGEM3.GC31.LL".

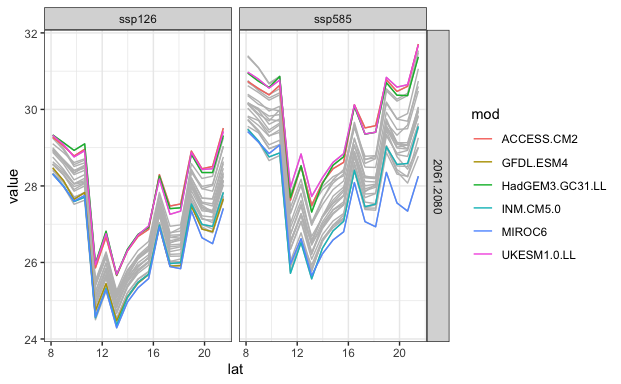

Supplementary Figure 1. Variation in mean annual temperature across the western ghats ridge across different climate models.

**Visualizing interaction between elevational extents and midpoints of Frogs**

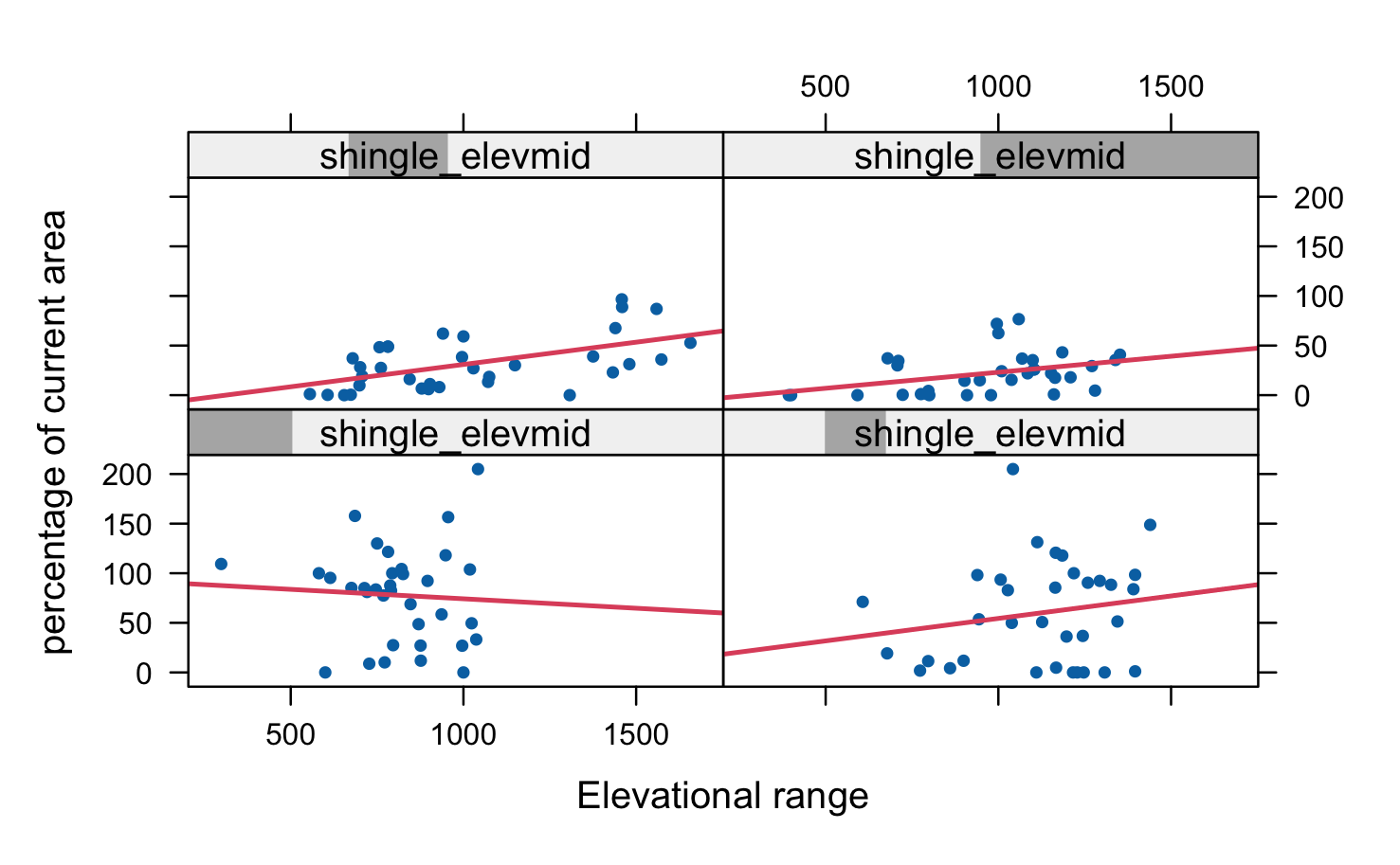

Supplementary figure 2. Predicted changes in suitable area across elevational ranges of species, conditional of elevational midpoints

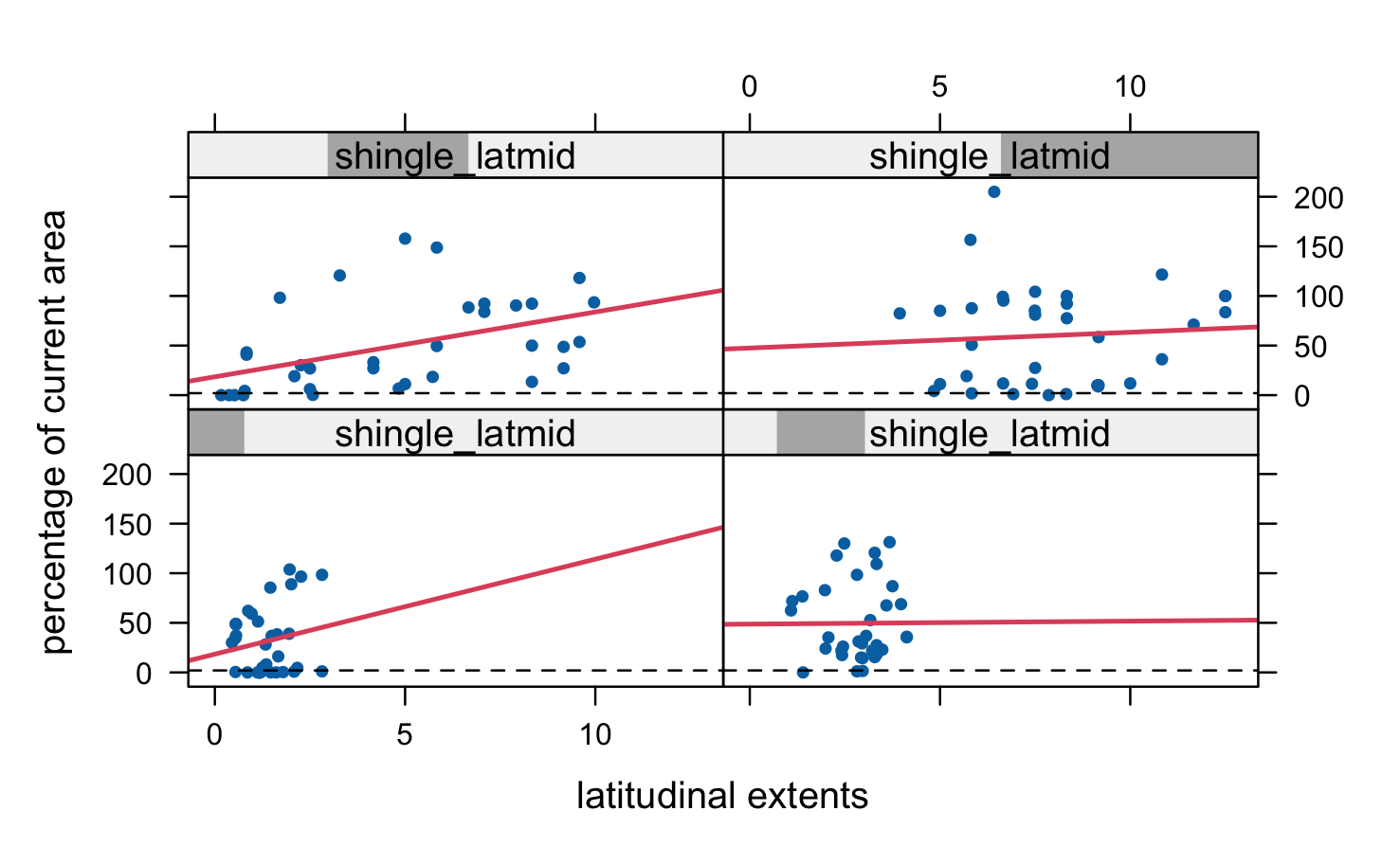

Supplementary figure 2. Predicted changes in suitable area across latitudinal ranges of species, conditional of latitudinal midpoints
